# Neurotropism of enterovirus D68 isolates is independent of sialic acid and is not a recently acquired phenotype

**DOI:** 10.1101/161778

**Authors:** Amy B Rosenfeld, Audrey L Warren, Vincent R Racaniello

## Abstract

Acute flaccid myelitis /acute flaccid paralysis (AFM/AFP) is a rare but serious illness of the nervous system, specifically affecting the grey matter of the spinal cord, motor controlling regions of the brain and the cranial nerve. Most cases of AFM/AFP are pathogen associated, typically with poliovirus and enterovirus infections, and occur in children under the age of 6 years old. Enterovirus D68 (EV-D68) was first isolated from children with pneumonia in 1962, but an association with AFM/AFP was not observed until the 2014 outbreak. Organotypic mouse brain slice cultures generated from postnatal day 1 to 10 mice were used to determine if neurotropism of EV-D68 is shared among virus isolates. Six of the seven EV-D68 isolates examined, including two from 1962 and four from the 2014 outbreak, replicated in neurons, and all replicated in astrocytes. Furthermore, a putative viral receptor, sialic acid, is not required for neurotropism of EV-D68, as both sialic acid dependent and independent viruses replicated within neurons. These observations demonstrate that EV-D68 is neurotropic independent of its genetic lineage, can infect both neurons and astrocytes, and that neurotropism is not a recently acquired characteristic as has been suggested.

**Significance:** Recently there has been an increase in the number of children infected with enterovirus D68 (EV-D68). Most infections are associated with mild flu-like symptoms, but neurological dysfunction may develop in a small number of children. How the biochemical and genetic differences among EV-D68 isolates relates to development of neurological disease remains an unanswered question. Assessing infection of multiple viral isolates in organotypic brain slice cultures from postnatal day 1 to 10 mice revealed that multiple isolates are neurotropic. Both neuraminidase sensitive and resistant viruses infected neurons, indicating that sialic acid binding does not play a role in EV-D68 neuropathogenesis. Establishment of a genetically and pharmacologically amenable system using organotypic brain slice cultures will provide insight into how EV-D68 neuropathologies develop.

## Introduction

The picornavirus enterovirus D68 (EV-D68) recently re-emerged as an infectious cause of severe respiratory distress in young children (1-12). Originally isolated from children with pneumonia in 1962 (13), few cases of EV-D68 infection were reported (https://www.cdc.gov/non-polio-enterovirus/about/ev-d68.html) until two outbreaks in the United States and Europe in the summer and fall of 2014 and June 2016 (1-12). During these epidemics numerous children diagnosed with severe respiratory distress induced by EV-D68 infection also developed an acute flaccid myelitis/paralysis (AFM/AFP) similar to that caused by poliovirus, a related enterovirus (2, 3, 6, 7, 10-12, 14). Phylogenetic analysis suggests that EV-D68 grouped within the B1 and B3 clades is associated with AFM/AFP (6, 15-19)(https://www.cdc.gov/acute-flaccid-myelitis/afm-surveillance.html). However, a 1962 virus isolate not within the B clade was found to be neurovirulent in suckling mice (13). Apart from two clinical samples, EV-D68 has not been isolated from the cerebrospinal fluid of infected children with AFM/AFP, nor has virus been found in the blood of these patients (20, 21).

EV-D68 is a unique enterovirus, as its physical and genetic properties are similar to both human rhinoviruses (HRVs) and poliovirus. Akin to the HRV particle, the EV-D68 virion is acid labile and optimal virus growth occurs at 33°C (13). Like HRVs, the sites of infection for EV-D68 are the nasopharyngeal cavity and the respiratory tract, but not the oropharyngeal and intestinal mucosa which are sites of entry for poliovirus.

Transmission of EV-D68 is also very different from that of poliovirus, and occurs via respiratory aerosols, not fecal-oral contamination (22). Yet EV-D68 disease is extremely different from that caused by either HRVs or polioviruses as EV-D68 infected patients do not develop a viremia, and virus is rarely shed in the feces of infected patients (22, 23).

Children with AFM/AFP associated with EV-D68 displayed limb weakness and cranial nerve abnormalities (6, 14, 15, 24, 25). It has been suggested that neurotropism of EV-D68 is a recently acquired phenotype (6, 26-28). To answer this question we used organotypic brain slice cultures from mice to examine the neurotropism of EV-D68 isolates from 1962 to 2014. As sialic acid binding varies among these isolates, it was also possible to assess the role of this sugar in neurotropism.

## Results

### EV-D68 infection of mouse neuroblastoma and embryonic fibroblasts

To determine if mouse cells are susceptible and permissive for EV-D68 infection, a mouse embryonic fibroblast cell line (MEF) and a mouse neuroblastoma cell line (N2A) were infected at a multiplicity of infection (MOI) of 3 with multiple isolates of EV-D68 including Fermon and Rhyne from the initial virus outbreak in 1962; NY from 2009, and 2014 isolates including 947, 949, 952, 953 and 956. Virus replication was assessed by plaque assay. The NY, 947 and Rhyne isolates replicated in MEFs, while all but the Rhyne isolate replicated in the N2A cultures (Fig. 1). These results demonstrate that some but not all isolates of EV-D68 can replicate in mouse cells.

**Figure 1.**
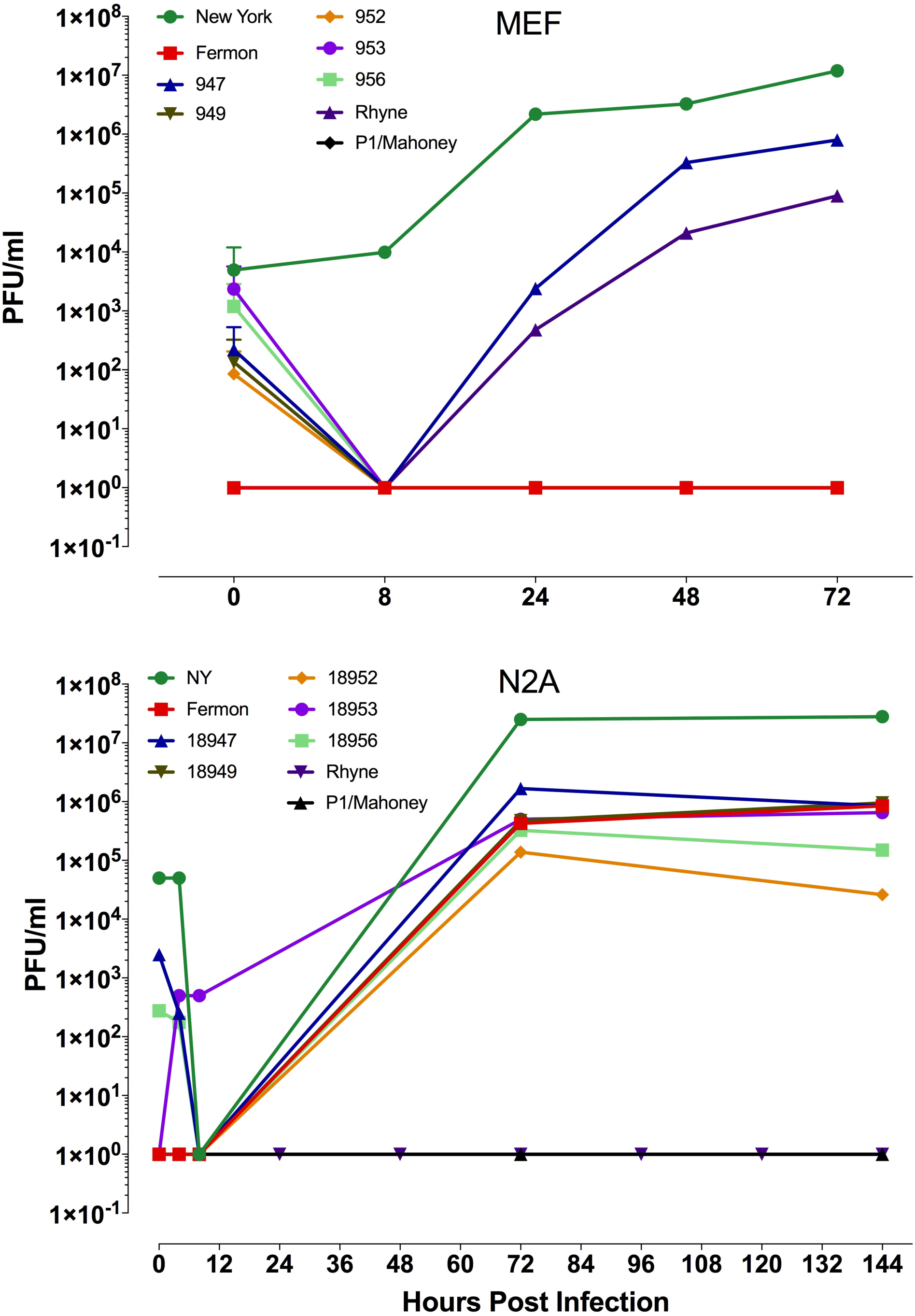
Replication of EV-D68 isolates in mouse embryonic fibroblasts and neuroblastoma cells. Time course of replication of multiple isolates of EV-D68 from mouse embryonic fibroblast cell line (MEF) (A) and a mouse neuroblastoma cell line (N2A) (B). Cells were infected with virus at an moi of 3 and incubated at 33°C, and at the indicated times cultures were harvested and assayed for infectious virus by plaque assay.

### Interaction of EV-D68 and sialic acid on the surface of human cells

Several reports, including the initial description of the virus, have shown that, *in vitro*, sialic acid binds the virus capsid (13, 29-32). Furthermore, the crystal structure of sialic acid bound to EV-D68 revealed that sialic acid binds within the canyon located at the 5-fold axis of symmetry (30, 33). To determine if sialic acid interacts with multiple isolates, hemagglutination assays were performed using red blood cells (RBCs) from both guinea pigs and humans. Nine different batches of human RBCs were used comprising 3 donors per batch. Fermon and three isolates from 2014 (947, 949, and 956) agglutinated guinea pig RBCs; no isolate agglutinated human RBCs (Table 1). The absence of human RBC agglutination was previously observed using EV-D68 isolates from the 2012 outbreak in the Netherlands (29).

**Table 1.** Hemagglutination of EV-D68 isolates on guinea pig and human red blood cells.

| Virus Isolate | Hemagglutination Titer |  |
| --- | --- | --- |
|  | Human | Guinea Pig |
| New York | 0 | 0 |
| Fermon | 0 | 256 |
| Rhyne | 0 | 0 |
| 947 | 0 | 256 |
| 949 | 0 | 64 |
| 952 | 0 | 0 |
| 953 | 0 | 0 |
| 956 | 0 | 512 |
| Influenza A/WSN | 256 | 1024 |

### Neuraminidase resistance is cell type and isolate specific

The requirement for sialic acid can also be investigated by removal of sialic acid from the cell surface with neuraminidase prior to virus infection. To further refine understanding of the requirement for sialic acid for infection with EV-D68, cultures of various cells including rhabdomyosarcoma (RD) cells, the cells used for propagation and titration of the virus, and N2A cells were incubated with medium containing neuraminidase 30 minutes prior to infection to remove sialic acid. Cells were infected at an MOI of 3, and at the indicated time post-infection viral replication was determined by plaque assay. All EV-D68 isolates were able to infect neuraminidase treated cells albeit with diminished efficiency (Fig. 2). The effect of neuraminidase treatment on viral yield varied between isolates and between cell types. These results suggest that interaction with sialic acid may be necessary in only certain contexts to facilitate virus attachment to the cell surface, as described for Theiler’s murine encephalomyelitis and enterovirus 70 (34-37).

**Figure 2.**
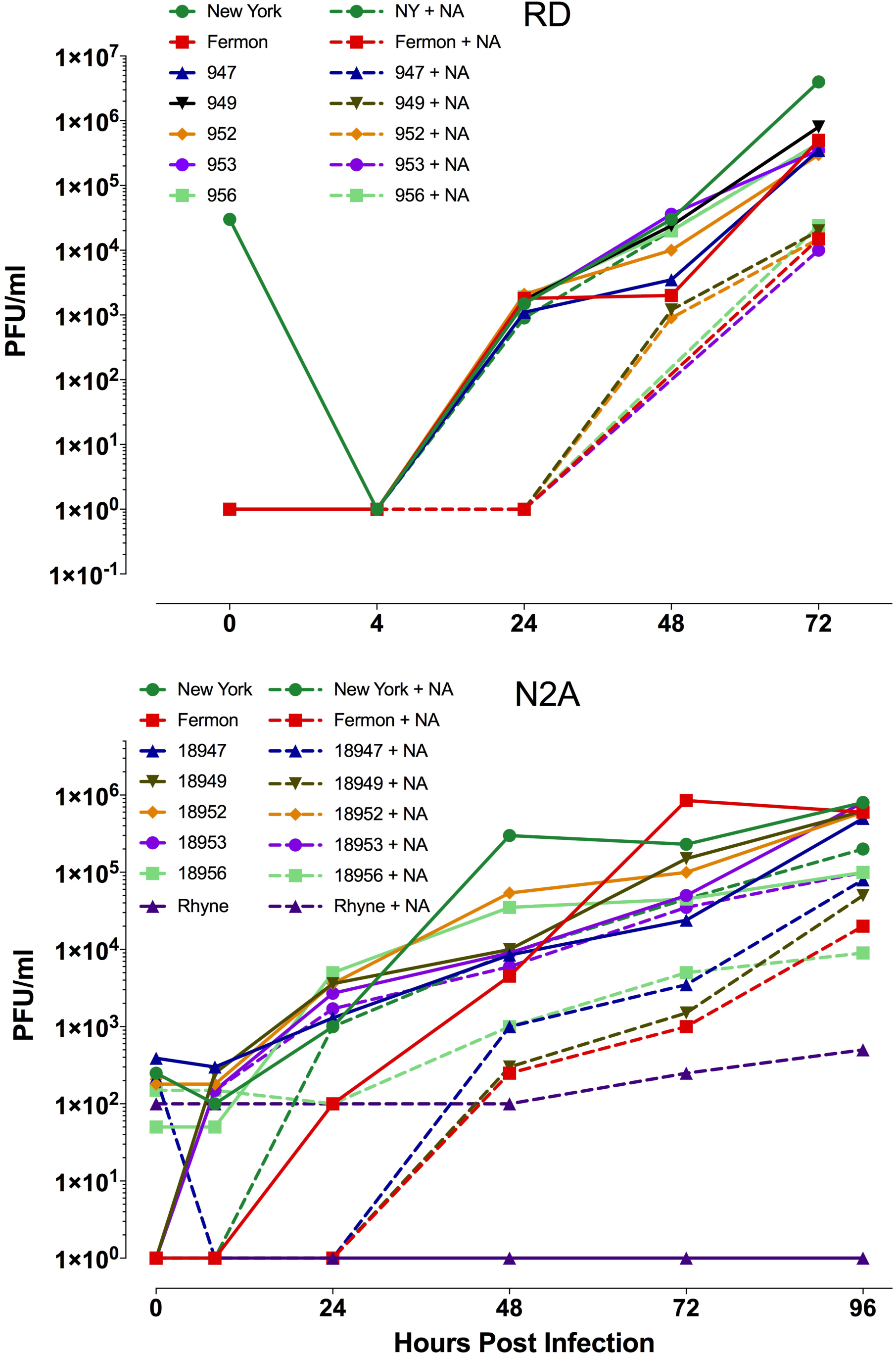
Effect of neuraminidase on replication of EV-D68 isolates. Time course of replication of multiple isolates of EV-D68 from RD (A) and N2A (B) cells untreated, or treated with neuraminidase at 37°C for 30 minutes prior to virus infection. Cells were infected with virus at an moi of 3 and incubated at 33°C, and at indicated times cultures were harvested and assayed for infectious virus by plaque assay.

### Replication of enterovirus D68 in post-natal organotypic brain slice cultures

Development of polio-like acute paralysis and neuronal dysfunction has been observed in children infected with EV-D68 (6, 15, 24). However, virus has been isolated from the CNS of only two infected children. While a murine model of EV-D68 induced paralytic myelitis has been established, this model only examines virus pathogenesis in one day old mice, and intramuscular infection of the virus is required for efficient development of paralysis (38). Unlike poliovirus, EV-D68 is a respiratory virus not known to infect the musculoskeletal system, nor has virus been observed to traffic into the CNS via the neuro-muscular junction. Furthermore, radiological images of infected patients have identified lesions within the brain stem specifically the cranial nerve motor nuclei (15, 25-27, 39, 40). Viral infection of brain was not examined in this model.

To determine if EV-D68 is neurotropic, organotypic brain slices generated from P2, P4 and P10 mice were infected with representative isolates. Virus replication was assessed by plaque assay at various times post infection. All EV-D68 isolates productively infected brain slice cultures from P2, P4, and P10 mice (Fig. 3). As expected, poliovirus did not replicate in murine brain slice cultures. Indirect immunofluorescence microscopy using antibodies to Iba1, a microglia specific calcium-binding protein, and the pan-enterovirus monoclonal antibody L66J as well as NeuroTrace Nissl stain, a dye that selectively binds Nissl substance found within in the cytoplasm of neurons within the brain and spinal cord, revealed that six EV-D68 isolates infected Nissl stained neurons (Fig. 4 and data not shown). Viral antigen was detected in brain slices infected with the 952 isolate, but it was not present in Nissl stained neurons (Fig. 4B) but most likely in astrocytes (data not shown).

**Figure 3.**
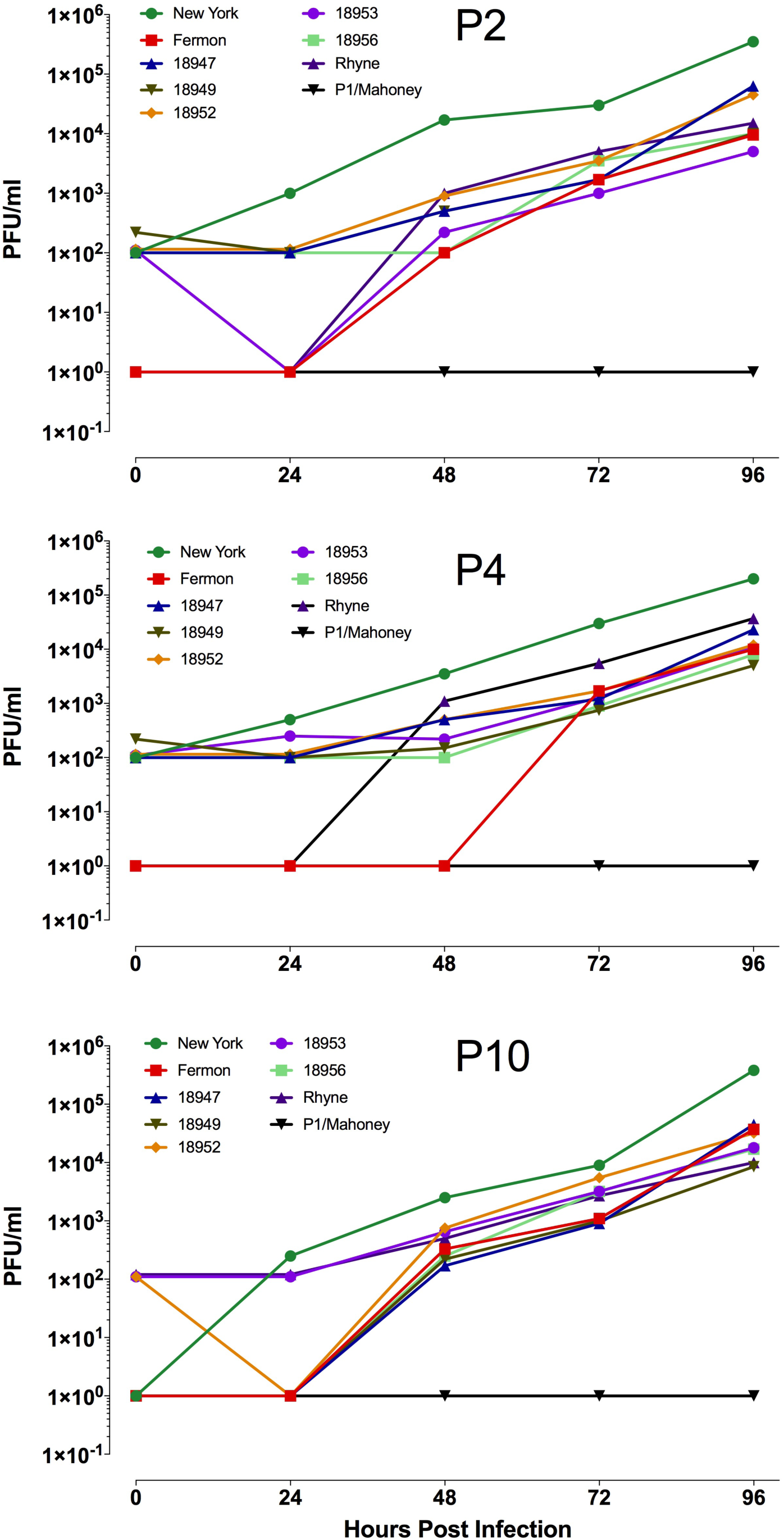
Replication of EV-D68 isolates in organotypic brain slice cultures generated from postnatal day 2, 4 and 10 mice. Time course of replication of multiple isolates of EV-D68 in organotypic brain slice cultures produced from postnatal day 2 (P2), 4 (P4) and 10 (P10) old mice. Cultures were infected with 105 pfu of virus and incubated at 33°C. At indicated times cultures were harvested and assayed for infectious virus by plaque assay.

**Figure 4.**
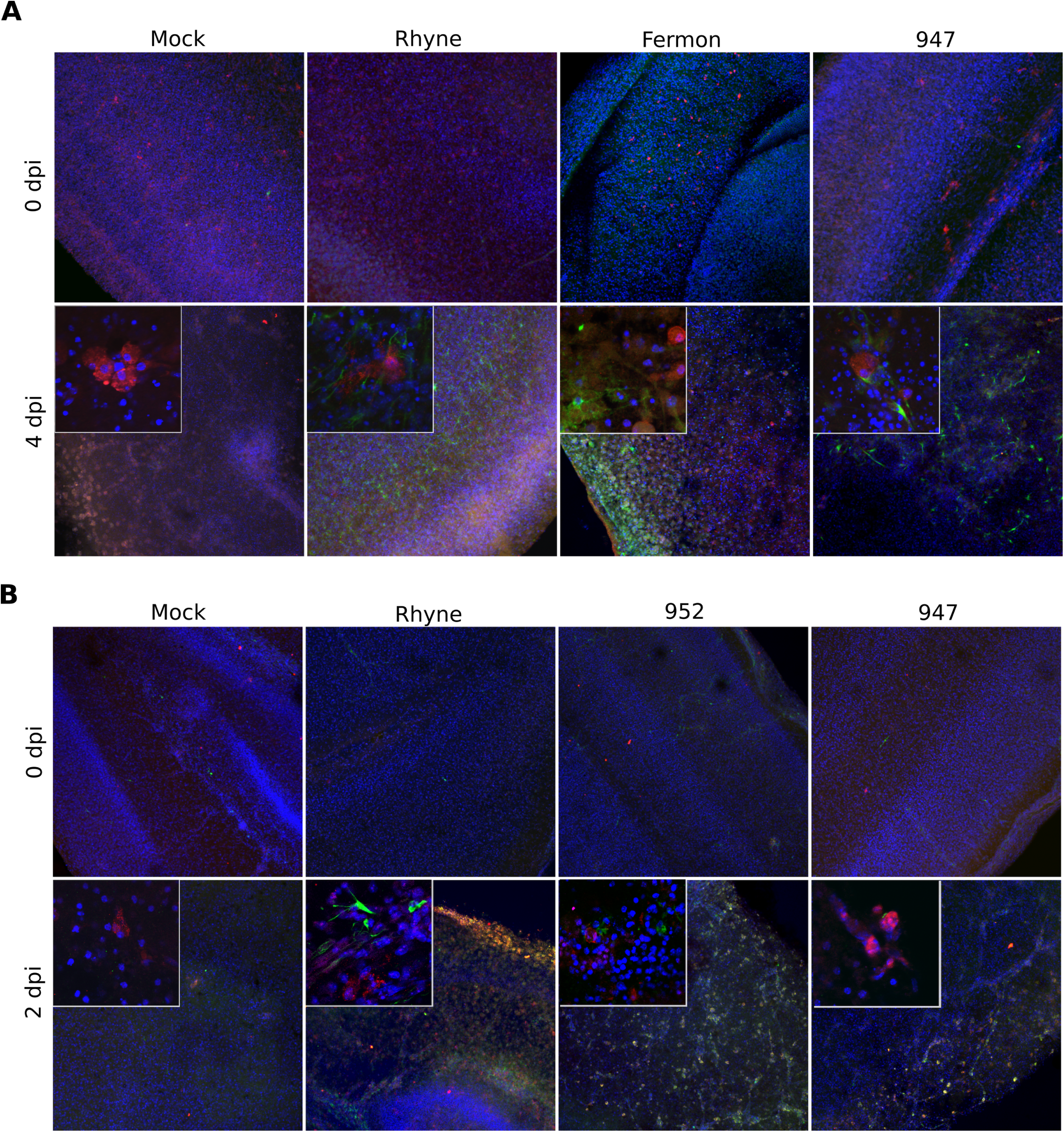
Indirect immunofluorescence microscopy of EV-D68 infected organotypic brain slice cultures. Indirect immunofluorescence microscopy of organotypic brain slice cultures generated from postnatal day 2 (A), and 10 (B) old mice infected with representative EV-D68 isolates. At indicated times cultures were fixed and stained with antibody to IBA-1(red), pan-enterovirus capsid antibody (green) and DAPI (blue).

To determine if the analogous cells of the human CNS also support virus replication, cortical neuron and astrocyte cultures derived from human induced pluripotent stem cells (hiPSCs) were infected with EV-D68 isolates. Sensory, inhibitory and excitatory neurons, along with motor neurons can be found within cortical neuron cultures. Culture medium was collected at various times post-infection and virus production was assessed by plaque assay. Replication of all isolates except 952 was observed in cultures of induced neurons (Fig. 5A). All seven isolates of EV-D68 replicated in human induced astrocyte cultures (Fig. 5B).

**Figure 5.**
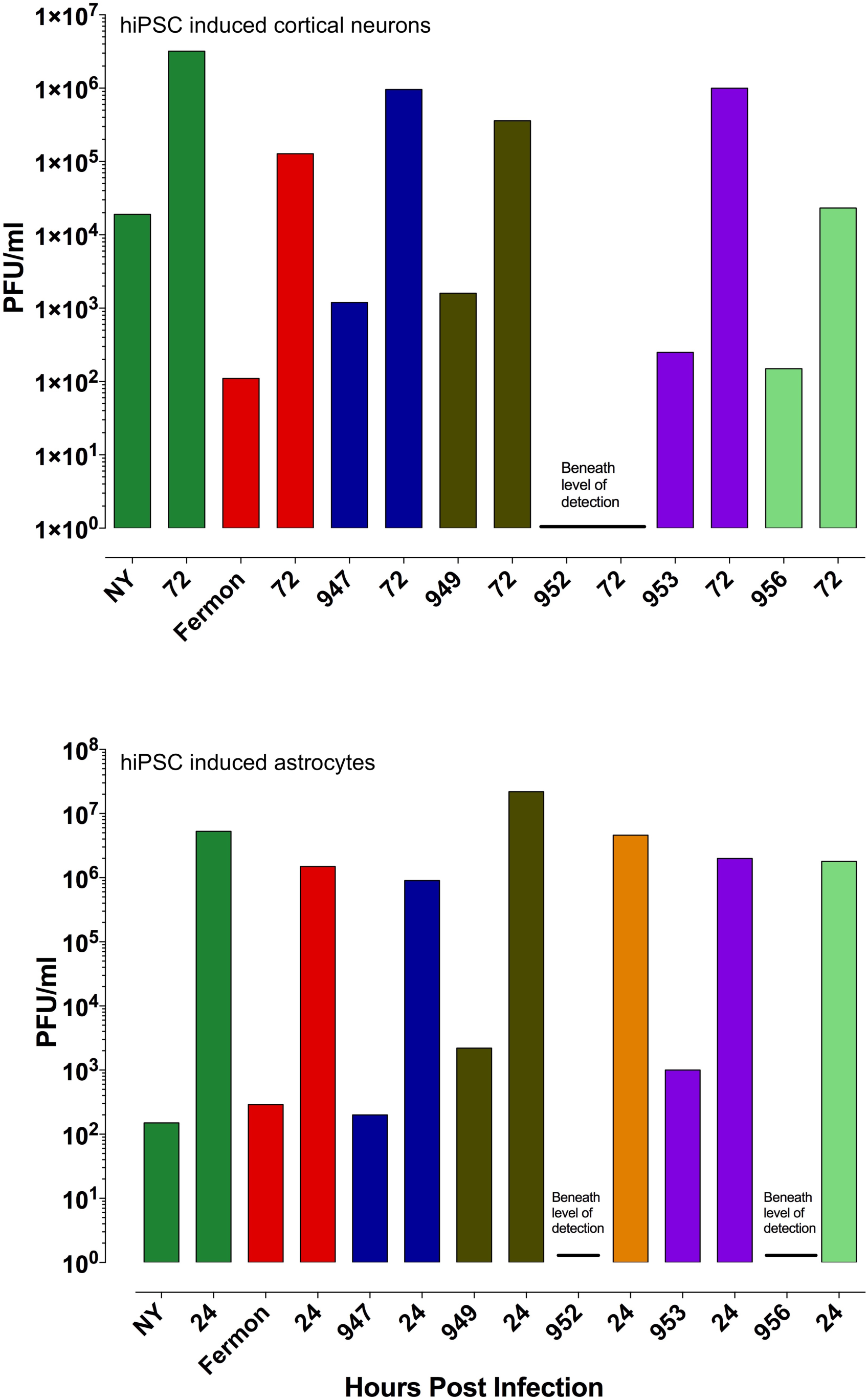
Replication of EV-D68 isolates in human cortical neurons derived from induced pluripotent stem cells. Time course of replication of multiple isolates of EV-D68 from cortical neurons (A) or astrocytes (B) derived from induced human pluripotent stem cells. Cells were infected with virus at an moi of 3 and incubated at 33°C, and at indicated times cultures were harvested and assayed for infectious virus by plaque assay.

## Discussion

Three animal models of EV-D68 infection have recently been established (38, 41, 42) but they are of limited utility for understanding the pathogenesis of neurological disease. In a cotton rat model, replication in the respiratory tract and the induction of neutralizing antibodies were studied after intranasal inoculation, but neurotropism and neurovirulence of EV-D68 were not investigated (41). A transient viremia develops in ferrets after intranasal inoculation, and virus specific RNA was detected in feces from infected animals, neither of which are observed during most human infections. No neurological disease developed in this animal model of EV-D68 infection (42). In a murine model of EV-D68 infection, in which postnatal day 1 mice were used, paralysis was observed after intracranial or intramuscular injection of the virus (38). In depth characterization of virus replication and induction of AFM/AFP was determined for only one EV-D68 isolate, which paralyzed approximately 50% of animals after intracranial inoculation. Furthermore, little to no neuronal dysfunction was observed when mice of the same age were intranasally infected, which is the natural route of infection in humans. In this mouse model, paralysis occurred most frequently when mice were intramuscularly injected, while the Rhyne strain of EV-D68 was not neurotropic after intracranial inoculation, in contrast to reports by others (13). While the authors suggest that EV-D68 may invade the CNS in a manner similar to poliovirus - via the neuromuscular junction – this hypothesis was not addressed, e.g. by severing the nerve that innervates the limb prior to intramuscular infection to impair trafficking. EV-D68 is a respiratory virus not known to infect the musculoskeletal system and rarely is a viremia established. The mechanism by which the virus traffics into the CNS is not known.

Organotypic brain slice cultures are ideal for studying EV-D68 neurotropism. Although production of brain slices causes severing nerve connections, organotypic brain slice cultures nevertheless maintain many aspects of *in vivo* brain biology, including functional local synaptic circuitry with preserved brain architecture, vascularization and immune composition, and can remain viable for several days in culture. Their utility has been well established in hundreds of publications. Organotypic cultures are highly reproducible because they are derived from exact stages of development. They have been extensively used to understand neuronal connectivity and neurodegenerative disorders, and are the experimental system for initial testing of viral vectors for gene therapy (43-47). They have also been used to study Zika virus infection (48). Organotypic brain slice cultures have enriched our understanding of autosomal-recessive microcephaly and brain development (49, 50). They not only complement findings from established animal models, but are easier to genetically and pharmacologically manipulate than live mice, and can be used to elucidate questions about cell biology. This culture system is also the only means by which neurotropism of a virus can be directly assessed independent of its ability to enter the nervous system, placing the virus inoculum directly on target cells. Intracranial infection does not fully separate neuroinvasion, the ability to traffic from the periphery into the CNS including crossing the blood brain barrier, from neurotropism, because virus is also delivered to the space surrounding the brain. Organoid cultures, while very popular, are unable to faithfully mimic the cellular and structural complexity of the brain, and are heterogeneous with respect to cell type composition and structure, which influences the reproducibility of any findings (51).

The results of phylogenetic analysis of EV-D68 isolates suggest that viruses from specific evolutionary lineages are associated with AFM/AFP (6, 26-28). We examined the ability of viruses from multiple lineages, including subclade B1, initially associated with the outbreak of AFM/AFP in 2014, for the ability to infect organotypic brain slice cultures from day 1 to 10 postnatal mice, developmental stages that correlate with the years 1-6 in human brain development. We find that all EV-D68 isolates examined can replicate in brain slice cultures. We conclude that EV-D68 is neurotropic independent of its genetic lineage, and that neurotropism is not a recently acquired characteristic as has been suggested (6, 26-28). Instead, our data align with recent phylogenetic analysis demonstrating that EV-D68 sequences from AFM/AFP patients do not form a monophyletic group but are interspersed among the clades (17, 38). The association of AFM/AFP with a specific genetic lineage may reflect the ability of certain isolates to invade the CNS. For example, effective modulation of innate immune defenses in the respiratory tract may promote neuroinvasion, and this property might be associated with specific genetic lineages.

It is intriguing that all but one isolate of EV-D68 that we examined can infect and replicate in neurons. An important question is whether similar results are observed in human cells. We find that all isolates replicate in human iPSC derived cortical neurons, except for isolate 952 (Fig. 5), which replicates, together with all other isolates, in human iPSC derived astrocytes (unpublished results). The 952 isolate was most efficient at inducing limb weakness in a murine model of EV-D68 pathogenesis (38). Presumably replication in neurons would lead to the symptoms of AFP, including muscle and cranial nerve dysfunction. How replication in astrocytes would lead to neuronal dysfunction and paralysis is not known. Replication of EV 71 in astrocytes is thought to participate in development of paralysis observed in virus infected humans, non-human primates, and mice (52). A role for infection of astrocytes has also been proposed to be part of the neuropathology caused by the flavivirus West Nile virus, and the alphavirus chikungunya (53, 54). The ability of West Nile virus to infect astrocytes is not only a distinguishing characteristic between pathogenic and non-pathogenic isolates, but leads to the release of neurotoxic factors responsible for neuronal apoptosis (55, 56). A role for astrocytosis in motor neuron destruction during amyotrophic lateral sclerosis has been proposed (57).

The identity of the receptor for EV-D68 is debatable. ICAM-5 has been suggested to function as a proteinaceous receptor for EV-D68 in the brain (32). However, production of ICAM-5 is restricted to the telencephalon, the most rostral segment of the brain region, consequently, it is not present on the surface of cells within the brain stem or spinal cord, two sites where lesions are found in radiological images of EV-D68 infected children (15, 25-27, 39, 40, 58). Furthermore, ICAM-5 is not found on the surface of epithelial cells that line the respiratory tract (http://www.proteinatlas.org/ENSG00000105376-ICAM5/tissue). The cells that we use to propagate EV-D68, including RD, N2A, and MEFs, do no produce cell surface ICAM-5 as determined by flow cytometry, nor do they produce ICAM-5 mRNA (data not shown).

The role of sialic acid in EV-D68 infection is unclear. Certain isolates of the virus can hemagglutinate red blood cells and can replicate in cells treated with neuraminidase to remove sialic acid (13, 29, 31, 33). This sugar binds the viral capsid on the eastern wall of the canyon located at the 5-fold axis of symmetry, displacing a C14 fatty acid from a hydrophobic pocket, allowing for the partial collapse of the canyon (30, 33). However, VP4 density was still observed in the structure of the EV-D68-sialic acid complex. Moreover no change in the size of the virion was observed in the presence of sialyated receptor analogues at room temperature, suggesting that sialic acid may only initiate the first steps of the entry process, attachment to the cell surface, and is not involved in conformational changes related to uncoating (33). Inactivation of the gene encoding cytidine monophosphate N-acetylneuraminic acid synthase, which is required for synthesis of sialic acids, impairs replication of some, but not all isolates of EV-D68 (29). However, it was not demonstrated that lack of sialic acid impairs virus binding to cells. Disruption of sialic acid production not only effects protein glycosylation, but also impairs synthesis of glycolipids. The formation and function of intracellular vesicles and proper fusion between these vesicles is dependent upon glycolipids. Such vesicles are critical for the entry and RNA synthesis of enteroviruses. Our results confirm that removal of sialic acid from the cell surface with neuraminidase impairs infection with specific isolates of EV-D68, but it is important to note that such treatment only allows for the function of extracellular sialic acid to be assessed. While the ability to hemagglutinate red blood cells varies according to isolate, all isolates tested replicate in organotypic brain slices from mice. We conclude that binding to sialic acid is not essential for infection of the brain.

The results presented here establish organotypic brain slice cultures as a pharmacologically and genetically amenable system to study EV-D68 neurotropism and neurovirulence that has distinct advantages compared with live animals and organoid cultures. For example, when assessing the effects of experimental drugs on viral replication in brain slice cultures, the added compounds are confined to the culture medium, rather than potentially altering the physiology of the entire animal. Slice cultures can be produced from mice of any genetic background, and DNA encoding mutant alleles can be readily introduced by *ex vivo* electroporation prior to generating cultures to study the effects of host protein variants on viral replication. The alternative, to knock-in alleles in mice, is far more time and reagent consuming. The use of brain slice cultures will provide mechanistic insight into EV-D68 neurotropism and neuropathogenesis.

## Materials and Methods

### Ethics Statement

All experiments were performed in accordance with guidelines from the Guide for the Care and Use of Laboratory Animals of the NIH. Protocols were reviewed and approved by the Institutional Animal Care and Use Committee (IACUC) at Columbia University School of Medicine (assurance number AC-AAAR5408).

### Cell and organotypic brain slice cultures from neonatal mice

Rhabdomyosarcoma (RD) cells were grown in Dulbecco’s modified Eagle medium (Invitrogen, Carlsbad, CA), 10% fetal calf serum (HyClone, Logan, UT), and 1% penicillin-streptomycin (Invitrogen).

C57/Black 6 mice were bred in a specific-pathogen-free facility at Columbia University Medical Center. At the appropriate time post birth, mice were euthanized per animal study protocol and brains from were dissected into ice-cold artificial cerebrospinal fluid (ACSF) consisting of 125mM NaCl, 5mM KCl, 1.25mM NaH_2_PO_4_, 1mM MgSO_4_, 2mM CaCl_2_, 25mM NaHCO_3_, and 20mM glucose, pH7.4, 310 mOsm1^-1^. Brains were embedded in 4% low melting point agarose dissolved in ACSF and sliced into 300ΰm coronal sections using a vibratome (Zeiss). Slices were maintained on 0.4ΰm, 30mm diameter Millicell-CM inserts (Millipore) in cortical culture medium (CCM) containing 25% Hanks balanced salt solution, 47% basal MEM, 25% normal horse serum, 1X penicillin-streptomycin-glutamine, and 30% glucose. Cultures were maintained in a humidified incubator at 37 °C with constant 5% CO2 supply.

### Viruses

Enterovirus D68 isolates (EV-D68): 18947 (, 18949 (, 18952 (, 18953 (and 18956 ( were obtained from BEI Resources. Rhyne and New York (NY) EV-D68 isolates were kindly provided by Shigeo Yagi (California Department of Public Health, Richmond, Ca) and W. Ian Lipkin (Mailman School of Public Health, Columbia University), respectively. The Fermon isolate was purchased from ATCC (American Tissue Culture Collection, Manassas, VA). All viruses were propagated and assayed in RD cells. Viral titers were determined by plaque assay (see below).

### Binding assays

Hemagglutination assays were done in V-shaped microtiter plates (Fisher, 07-200-698). Equal volumes of 0.5% washed human (David Fidock, Columbia University, NY) or guinea pig (Rockland, R402-0050) red blood cells were added to serial 2-fold dilutions of virus. Plates were incubated at 4°C and readings were made every 30 minutes for 2 hours. Activity of neuraminidase was verified by lack of influenza virus hemagglutination on treated guinea pig red blood cells (data not shown).

### Indirect immunofluorescence microscopy

At either 0 h or 96h post infection the medium was removed from EV-D68 virus infected or uninfected organotypic brain slices cultures and the cultures were placed overnight in 4% paraformaldehyde (PFA) fixative dissolved in 1X PBS at 4 °C. Following fixation, cultures were incubated in blocking solution of PBS, 0.3% Triton X-100 and 3% horse serum. Cultures were incubated overnight in at 4 °C in blocking solution containing appropriate primary antibodies. Sections were washed in 1X PBS, and incubated in the presence of fluorophore-conjugated secondary in blocking solution. Sections were mounted on slides using Aqua-Poly/Mount (Polysciences, Inc) or Fluorsave (Millipore, Fisher) and imaged using a iZ80 laser scanning confocal microscope (Olympus FV100 spectral confocal system). Brain sections were imaged using a X60 1.42 N>A> oil objective or a X 10 0.40 N.A. air objective. All images were analyzed using ImageJ software (NIH, Bethesda, MD, USA).

### Plaque assay

RD cells were seeded on 60mm plates for approximately 70% confluence at the time of plaquing. Next, 100μL portions of serial 10-fold virus dilutions were incubated with cells for 1 h at 37 °C. Two overlays were added to the infected cells. The first overlay consisted of 2 ml of 1 DMEM, 0.8% Noble agar, 0.1% bovine serum albumin, 40 mM MgCl_2_, and 10% bovine calf serum. After solidification, a second liquid overlay was added that was composed of 1 DMEM, 0.1% bovine serum albumin, 40 mM MgCl_2_, 0.2% glucose, 2 mM pyruvate, 4 mM glutamine, and 4 mM oxaloacetic acid. The cells were incubated at 37 °C for 4-6 days and developed by using 10% trichloroacetic acid and crystal violet.

### Virus infections

Organotypic brain slice cultures were infected with 105 pfu of EV-D68 isolates: NY, Fermon, Rhyne, 947, 949, 952, 953, 956 or poliovirus type 1. Virus was allowed to adsorb to the slices for 1h at 37 °C. The inoculum was removed, and the slices were washed 2X in 1X PBS. Infected slices were cultured in CCM for 96 hours.

### Antibodies

Antibodies used in this study were mouse monoclonal against capsid proteins (LifeTechnologies, MA518206, 1:250 dilution), rabbit polyclonal against glial fibrillary acidic protein (GFAP) (Abcam, ab53554), and rabbit polyclonal allograft inflammatory factor (Iba1) (Wako, 019-19741). Donkey fluorophore-conjugated secondary antibodies (Jackson Labs, 1:500 dilution) were used together with DAPI (4’,6-diamidino-2-phenylindole, Thermo Scientific, 62248, 1:1,000 dilution) and NeuroTrace Nissl stain (LifeTechnologies, N21483).

### Data Analysis

GraphPad Prism software was used to analyze all data. Log10-transformed titers were used for graphing the results of plaque assays.

## Acknowledgments

This work was supported by the National Institutes of Health (grant AI121944).

We thank Richard Vallee for the use of laboratory equipment for producing brain slice cultures and for immunofluorescent analysis, and Ian Lipkin (Mailman School of Public Health, Columbia University) and Shigeo Yagi (California Department of Public Health, Richmond, Ca) for EV-D68 isolates.

